# Discrimination of non-nestmate early brood in ants: behavioural and chemical analyses

**DOI:** 10.1101/2021.03.17.435807

**Authors:** Arthur de Fouchier, Chloé Leroy, Abderrahman Khila, Patrizia d’Ettorre

**Author notes:** Corresponding author: Arthur de Fouchier, IEES-Paris, Campus Pierre et Marie Curie, 4 place Jussieu, 75005 Paris FRANCE.

## Abstract

Brood is critically important in social insect colonies. It carries the colony’s fitness through delivering future reproductive adults as well as workers that will increase the colony’s workforce. Adoption of non-nestmate brood can increase the colony workforce but entails the risk of rearing unrelated sexuals or social parasites. Thus, theory would predict that ant workers will evolve the ability to discriminate between nestmate and alien brood using the chemical cues displayed at the brood’s surface. This appears especially true for eggs and first instar (L1) larvae, which require more resources before becoming adult workers compared to older brood. However, the chemical signature of ant early brood stages and its recognition by workers remains understudied. To fill this gap, we investigated the chemical basis of early brood nestmate and cross-species recognition in six ant species. We also tested the discrimination behaviour of workers in brood retrieval trials. We observed species-level cues and discrimination against hetero-specific brood. We also found that eggs and most L1 larvae displayed a colony signature. However, only some species discriminated against non-nestmate early brood. Interestingly, these species belong to genera subject to brood parasitism. We hypothesize that non-nestmate brood discrimination could arise from species adaptations against brood parasitism.

## Introduction

Recognizing offspring is a key issue for parents in many animal species. It allows them to increase their fitness through proper investment in parental care (Trivers, 1972). Parents use various cues to recognize their kin, including acoustic (Searby et al., 2004), visual (Mateo, 2015), chemosensory (d’Ettorre, 2020) or contextual cues (Penn & Frommen, 2010). A well-known example of failed kin recognition that leads to decreased fitness is the cuckoo bird brood parasitism (Payne & Sorensen, 2005). Cuckoo birds take advantage of parents that care for chicks that hatch in their nest (contextual cue) and lay their eggs in the nest of these host birds. The cuckoo chick typically hatches first, discards other eggs in the nest, and becomes the only recipient of care from the host parents.

Hymenopteran social insects (some wasps and bees, and all the ants) are classical models for kin recognition as well (d’Ettorre, 2020). They usually recognise nestmates, that is individuals from the same group (colony), as a proxy of kin recognition (Bos & d’Ettorre, 2012). This recognition is important for cooperating with nestmates while competing for resources with non-nestmates. Kin and/or nestmate recognition is even more important for social insects, compared to many other social animals, as they show reproductive division of labour. This means that workers, which are fully or virtually sterile (Fletcher & Ross, 1985; Khila & Abouheif, 2008, 2010), achieve fitness indirectly by rearing their mother’s brood. This provides future reproductive individuals (males and queens) or increases the workforce of colony to ultimately produce more offspring.

This reproductive division of labour is a hallmark of highly social societies and places brood at the centre of ant colonies. Workers promptly retrieve eggs and larvae found outside the nest (Lenoir, 1981), and secure them in case of colony disturbance (Meudec, 1978). Behavioural studies have shown that ant workers adopt brood from other nests, and even other species, while keeping a preference for nestmate eggs and larvae (Schultner & Pulliainen, 2020).

Brood adoption is an adaptive behaviour as larvae raised in a foreign and unrelated nest may eventually integrate the colony’s workforce (Fénéron & Jaisson, 1995; Fouks et al., 2011). Incipient colonies of *Lasius niger* and *Messor pergandei* often raid brood from neighbouring colonies to increase their chance of survival (Madsen & Offenberg, 2017). Brood theft can also take place during nest relocation (Paul & Annagiri, 2019). However, adopting non-nestmate brood entails a risk. Some ant species are subject to social parasites, which take advantage of the workers of the host colony to raise their own brood, which has a negative impact on the fitness of host colonies (Buschinger, 2009; Lenoir et al., 2001).

In theory, adopting non-nestmate brood involves a trade-off, for ant workers, between the gain of future workforce and the potential cost of raising unrelated reproductive individuals or a social parasite (Fouks et al., 2011). It appears thus adaptive to develop counter-measures to avoid such risks. The net gain of adopting early brood, eggs and first instar (L1) larvae, is decreased by the higher amount of resources needed for such brood to develop into workers. Furthermore, early female brood caste is usually not yet determined (Trible & Kronauer, 2017), which further increases the risk of adopting an unrelated future queen. Among the possible adaptations, there is the ability of workers to recognize intruding non-nestmate adults and brood (Satoi & Iwasa, 2019). Stricter discrimination against individuals not matching the colony’s signature entails a risk of recognition errors (Reeve, 1989), but appears beneficial when it occurs in populations subject to brood parasitism (Grüter et al., 2018). While one could predict that parasitized species could develop such adaptation, the accuracy of this hypothesis remains elusive (Buschinger, 2009; Lenoir et al., 2001).

Ants are usually efficient in recognizing non-nestmates and behave aggressively toward competitors (Sturgis & Gordon, 2012). Nestmate recognition relies on the detection of colony-specific chemosensory cues. These are mostly long chain hydrocarbons found on the outer surface of developing and adult individuals. The hydrocarbons can be linear and saturated (*n*-alkanes), unsaturated (alkenes), or contain methyl groups (methyl-branched alkanes) (van Zweden et al., 2010; van Zweden & d’Ettorre, 2010). The blend of hydrocarbons displayed by each individual is the result of both genetic (*e.g*., van Zweden et al., 2010) and environmental factors (*e.g*., Liang & Silverman, 2000). Cuticular cues homogenise between members of the colony through mutual grooming, food sharing (trophallaxis), inter-individual contacts or contact with the nest-material (Lenoir et al., 2009; van Zweden et al., 2010). Consequently, members of the same colony, which are typically closely related and live in the same environment, share similar cuticular chemical profiles.

The interest in brood nestmate recognition behaviour in ant colonies led to, at least, 40 studies in 33 ant species (brood recognition has been recently reviewed in Schultner & Pulliainen, 2020). However, these studies focused mostly on mid to late-stage larvae. Hydrocarbons displayed on ant eggs have been studied in few genera (d’Ettorre et al., 2004; Endler et al., 2004; Helanterä & d’Ettorre, 2015; Holman et al., 2010; Ruel et al., 2013; Tannure-Nascimento et al., 2009; van Zweden et al., 2009). To our knowledge, a colony-level signature of the surface hydrocarbons of the eggs has been convincingly found in two genera, belonging to the Ponerinae and the Formicinae (Helanterä & d’Ettorre, 2015; Tannure-Nascimento et al., 2009). Therefore, further studying the chemical signatures on eggs is necessary to better understand if and how they can be recognised as nestmate brood.

Brood can acquire the hydrocarbon signature through various mechanisms. The source of colony-level cues on brood is better known in eggs than in larvae. Freshly deposited eggs already bear the colony signature (Helanterä & d’Ettorre, 2015). Mothers appear to deposit hydrocarbons on eggs while they are maturing in their ovaries (Endler et al., 2004). Once laid, the surface hydrocarbons of the eggs could be influenced by contact with workers and allo-grooming (Schultner et al., 2017; van Zweden et al., 2010). However, the effect of contact alone is probably not a rapid process (d’Ettorre et al., 2006), and thus it might not be impactful, given the short duration of the early brood stages (a few days). It is possible that embryos produce hydrocarbons that might traverse the chorion through pores and modify the egg surface hydrocarbons (Juárez & Fernández, 2007).

Surface hydrocarbons and nestmate recognition of early stage larvae remains critically understudied. When larvae hatch from their egg, it is unclear if the egg surface hydrocarbons are transferred to the larvae or if freshly hatched larvae shall *de novo* synthesize their surface hydrocarbons (Howard & Blomquist, 2004). In the ant *Aphaenogaster senilis*, the amount of surface hydrocarbons on larvae is smaller compared to eggs and workers (Villalta et al., 2016). It is likely that most of the hydrocarbons on the surface of eggs are not transferred to the larvae. As such, whether first instar larvae display enough cues to be recognised as nestmates remains an open question.

In this study, we aimed at filling the gap in our knowledge of nestmate recognition of early brood stages in ants. We investigated the colony-level signature of surface hydrocarbons of eggs and first instar (L1) larvae from six species belonging to three different subfamilies: Myrmicinae, Formicinae and Dolichoderinae. To assess how selective workers are when adopting brood, we studied brood-oriented behaviour of workers facing eggs and L1 larvae originating from their colony (nestmate), from another homo-specific colony (non-nestmate) or from another species (hetero-specific).

## Material and methods

### Ant colonies collection and rearing

We used colonies of six ant species: *A. senilis* (Formicidae, Myrmicinae), *Camponotus aethiops* (Formicidae, Formicinae), *Formica fusca* (Formicidae, Formicinae), *L. niger* (Formicidae, Formicinae), *Messor barbarus* (Formicidae, Myrmicinae) and *Tapinoma darioi* (Formicidae, Dolichoderinae). The geographic distribution of all species pairs studied here, for instance *L. niger* and *F. fusca*, are partially overlapping (see https://antarea.fr/). We observed that some of the colonies of *A. senilis* collected lived with neighbouring *M. barbarus* colonies. The same for *F. fusca* colonies and *L. niger* colonies.

The *T. darioi* colonies were collected in October 2018 and February 2020 in the region of Argelès-sur-mer (France). The *A. senilis* colonies were collected around Argelès-sur-mer (France) in October 2018 and in the Doñana National Park (Spain) in March 2019. The *C. aethiops* colonies were collected in 2014 and 2016 around Toulouse (France). *L. niger* colonies originated from founding queens collected in 2018 in the region of Paris (France). *M. barbarus* colonies originated from founding queens collected in October 2017 in Saint Gilles (France). *F. fusca* colonies were collected in 2017 and 2019 in the Ermenonville forest (France). All ants were housed in artificial nests with plaster floor placed in a larger plastic box constituting the foraging arena. Colonies were kept under controlled laboratory conditions (25±2°C, 50±10% relative humidity, 12 h/12 h: day/night) and fed twice a week with dead crickets and a mixture of honey and apples. Water was provided *ad libitum*. Behavioural experiments were performed in 2019 and 2020. All experiments were performed after at least 1 month of laboratory rearing.

### Chemical analyses

Chemical analyses were performed in 2019 and 2020. Ant colonies were reared at least 3 months in laboratory conditions before the chemical analyses. We collected live eggs and larvae from nest-boxes of the six ant species. To obtain L1 larvae, we selected those of a size comparable to an egg that we found among egg piles. Despite the similar size between L1 larvae when they are folded on themselves and eggs, which is consistent with L1 larvae having hatched from an egg a few hours earlier, L1 larvae appear longer than egg but less large.

The number of eggs and larvae collected for chemical analyses is shown in Table S1. We collected at least three eggs and first instar (L1) larvae from at least three different colonies for each of the six species. Eggs and larvae were put individually into glass vial with a 200-µL glass insert (Supelco, Sigma-Aldrich) and immediately frozen. Surface chemicals extraction and analysis were performed within 6 months. Surface hydrocarbons were extracted from individual eggs and larvae using 10µl of n-pentane (≥99%, HPLC grade, Sigma-Aldrich) for 2 minutes. We then injected 3 µL of the extract into an Agilent 7890A gas chromatograph (GC), equipped with an HP-5MS capillary column (30 m x 0.25 mm x 0.25 µm) and a split-splitless injector, coupled to an Agilent 5975C mass spectrometer (MS) with 70 eV electron impact ionization. The carrier gas was helium at 1 mL.min^-1^. The temperature program was as follows: an initial hold at 70°C for 1 min, then 70-180°C at 30°C.min^-1^, then 180-300°C at 3°C.min^-1^, then 300-320°C at 20°C.min^-1^ then hold at 320°C for 3 min.

In order to assess the variations in the total amount of cuticular hydrocarbons between eggs and L1 larvae across species, we extracted additional samples from some of the species studied, depending on availability at the time of this experiment. The samples were collected and analysed by GC-MS as above except we added an internal standard in the solvent (pentane) used for the extraction (*n*-C_20_at 0,25ng/µL). The quantity of the surface hydrocarbons in the samples could then be estimated based on the area of this internal standard peak.

### Behavioural experiments

The aim was to test the behaviour of workers when facing nestmate, homo-specific non-nestmate or hetero-specific eggs or first instar larvae. The same protocol was followed for eggs and L1 larvae trials, which were performed independently. Overall, for the behavioural experiments, eggs and L1 larvae and workers originated from twelve *A. senilis* colonies, ten *C. aethiops, L. niger* and *M. barbarus* colonies and six *T. darioi* and *F. fusca* colonies. We prepared groups of six nestmate workers: three from outside the nest and three from inside the nest. This choice aimed at representing the diversity of age and role among workers in a colony, as workers found outside the nest tend to be older as well as foragers and workers from inside the nest tend to be younger and nurses. The ants were placed in an eight cm arena with a filter paper as floor and with walls coated with Fluon® (AGC Chemicals Europe, Thornton-Cleveleys, United Kingdom). Each group was given a refuge made of a red-coated 1.5mL Eppendorf tube (that had spent at least twenty-four hours in the nest box of the original colony), three late-instar larvae from their own colony, food (mixture of honey and apple) and water. After minimum time of twenty-six hours of acclimation, and if the workers had brought the late-instar larvae into the refuge, we removed food and water and started the behavioural trials. Groups that did not bring larvae into the refuge were discarded.

Shortly before the trials, we collected eggs or L1 larvae from the colony of origin of each group of tested workers (nestmate), from another colony of the same species (non-nestmate) or from another species (hetero-specific). For hetero-specific brood, we used brood from species of the same subfamily when possible to reduce the impact of the phylogenetic distance in recognition. We also choose brood from species of a similar size to reduce the impact of this cue in recognition. For *A. senilis*, we used *M. barbarus* brood and *vice versa*. For *C. aethiops* and *L. niger*, we used *F. fusca* brood. For *T. darioi*, we used *L. niger* brood. For each trial, three brood items were deposited in a line (figure A1). All three of these were either nestmate, or non-nestmate or hetero-specific relative to the workers. The behaviour of the workers towards the brood items was video recorded with an FDR-AX33 Sony camera for fifteen minutes. After fifteen additional minutes, any brood that had not been brought inside the refuge were removed from the arena. Thirty minutes later, another set of three brood items with a different origin were presented to the same group of workers. Each group of workers received nine brood items in total (all the three possible origins) in three different trials, in each trial the brood had the same origin. The different order of presentation of the three types of brood items were tested in an equilibrated manner between groups to prevent any bias. That is some groups received nestmate then non-nestmate then hetero-specific brood and an equivalent number of groups received nestmate then hetero-specific then non-nestmate, *etc*.).

The behaviour of the workers was scored for the first 15 minutes after the first brood item was deposited using the software Boris v7.9.15 (Friard & Gamba, 2016). We noted the times where workers started and stopped to antennate a brood item and the times when a worker entered the refuge with a transported brood item. The occurrences of aggressive behaviours (e.g., workers opening their mandibles, thus showing threat behaviour) towards homo-specific non-nestmate brood were very rare therefore we did not analyse such behaviours. Trials for which the workers did not touched or interacted with the brood items were discarded from further analysis as workers were considered inactive. Full details on the colonies and the number of groups used for each colony are displayed in supplementary table S1. For eggs, we used 36 groups from 6 *A. senilis* colonies; 39 groups from 4 *C. aethiops* colonies; 52 groups from 7 *L. niger* colonies; 36 groups from 6 *M. barbarus* colonies and 36 groups from 3 *T. darioi* colonies. For L1 larvae, we used 31 groups from 6 *A. senilis* colonies; 32 groups from 6 *C. aethiops* colonies; 40 groups from 7 *L. niger* colonies; 36 groups from 8 *M. barbarus* colonies and 32 groups from 3 *T. darioi* colonies. All experiments and scoring were performed by A. de Fouchier, except for *A. senilis* and *C. aethiops* L1 larvae experiments and for the scoring of *M. barbarus* eggs experiments that were performed under A. de Fouchier close supervision by two Master students.

### Data and statistical analyses

Data was analysed using R Studio (v1.3.1093, RStudio Team, 2015) and R software (v4.0.0, R Core Team, 2020). Data and code used for the analysis performed have been deposited on FigShare (doi: 10.6084/m9.figshare.14303078 and 10.6084/m9.figshare.14304167).

#### Chemical data

For each colony and species, we analysed between three and four samples (supplementary table S1). Hydrocarbons were identified by their mass spectra and retention times. Their areas were integrated using MSD ChemStation (vE.02.01.1177, Agilent Technologies Inc., CA), this was performed by A. de Fouchier.

The area of each peak was normalised to the sum of the area of all peaks in a given sample. To assess the variability of the chemical profiles across species and sample types, we performed a non-metric multidimensional scaling on the normalised areas of the peaks observed in all samples. This scaling was performed with three dimensions to give a good representation of the raw data (stress inferior at 0.1) and with 100 iteration maximum using the *metaMDS* function from the *vegan* package (v2.5-7).

For further analysis, we selected peaks that were present in all the samples of the same species. For the egg samples, the number of peaks was 18 for *A. senilis*, 28 for *C. aethiops*, 23 for *F. fusca*, 21 for *L. niger*, 16 for *M. barbarus* and 22 for *T. darioi*. For the L1 larvae samples, the number of peaks was 8 for *A. senilis*, 9 for *C. aethiops*, 5 for *F. fusca*, 5 for *L. niger*, 7 for *M. barbarus* and 6 for *T. darioi*.

We then did a principal component analysis (PCA) for each species using the *PCA* function (*FactoMineR* package, v2.0; Lê et al., 2008) and kept enough components to describe at least 95% of the total variance. We selected as subset of components an F-score, relative to the colony of origin, superior or equal to 0.01. The F-scores were computed with the function *fscore* (*PredPsych* package v0.4, Koul et al., 2018). Using those selected components, we computed linear discriminant analysis using the *LinearDA* function for each species and brood types separately using the colony of origin as classification variable with a leave-one sample out cross-validation (*PredPsych* package v0.4). To test the significance of the accuracy of classification obtained, we used permutation tests with 5000 simulations using the *ClassPerm* function (*PredPsych* package v0.4). This tests if the classification is more accurate than would be a random classification. This analysis was replicated using a different method to reduce complexity of the original dataset. We used dimensions from a non-metric multidimensional scaling on the normalised area of the peaks observed in all samples from the same species and sample type. This scaling was performed with the same tool as above but with enough dimensions to obtain a stress inferior at 0.05.

To assess the variability of the difference between nestmate and non-nestmate chemical signatures across species, we used the same datasets to compute intra and inter-colony Euclidean distance between nestmates and non-nestmates using the global centroid method (van Zweden et al., 2014). That is intra-colony distances are measured between each individual sample profiles and the mean profile of the colony. The inter-colony distances are measured between individual sample profiles and the mean profile of the samples from both the colony of origin of the individual sample under scrutiny and another colony. This allows to consider the variability between nestmate when measuring the distances with non-nestmates. In order to assess the variation of intra-colony distances between species, we computed the ratio between intra and inter-colony distances. That is, we normalised the intra-colony distances measures for each individual by dividing them by each inter-colony distances measured for the same individual. To assess the variation of intra-colony distances between species, we computed the ratio between intra and inter-colony distances. We then performed type II ANOVA, using the *Anova* function (*car* package, v3.0-7), on linear mixed-effects models (LMM), using the *lmer* function (*lme4* package, v1.1-23). The models were computed to test for the effect of the species of origin of the samples on a base 10 logarithmic transformation of the ratios of the intra and inter-colony chemical distances. Sample ID and colony of origin were used as nested random factors. The colony used for the inter-colony distance was a random factor as well. *P* values were adjusted for multiple comparisons across species for each type of brood using Holm’s method using the *p.adjust* function (package stats v4.0.0).

#### Behavioural data

We tested whether the source of the brood item had an effect on two different variables: 1) the number brood items brought into the refuge in each trial; 2) the total time workers spent antennating the brood items. The percentage of brood items brought to refuge was analysed using generalized linear mixed-effect models (GLMM) for proportional data with a binomial function with a logit link using the *glmer* function (package *lme4* v1.1-21). For the cumulative duration of antennation, we used LMMs using the *lmer* function (package *lme4* v1.1-21). The colony of origin of the workers, their group identity, the origin and the order of the brood encountered during the three trials were used as random factors for both types of models. Post hoc differences were tested with type II ANOVAs as above. *P* values were adjusted for multiple comparisons as above.

### Ethical Note

No licences or permits are needed for experiments on ants in France. We used 2220 adult worker ants for our behavioural experiments. We used 69 eggs and 74 L1 larvae for our chemical analyses. To minimise stress induced by rearing conditions, we used artificial nests with suiting humidity and foraging areas. Colonies were kept at optimal temperature and provided with sufficient food and water. No adult ants were disposed of during or after the experiment. Colonies for which the queen died after the experiments were disposed of by putting them at −20°C for at least 24h. Eggs and L1 larvae were sacrificed in a similar manner before solvent chemical extraction. No potentially harmful or painful manipulations of live animals were performed. No invasive samples were taken from live animals.

## Results

### Brood surface hydrocarbons

A non-metric multidimensional scaling ordination of the chemical profiles observed across samples, from all species and both types of brood, reveals that there is a clear difference between the profiles of all these categories (figure 1). This inter-specific and between brood type difference can be observed in the qualitative and quantitative variations of the hydrocarbons found in the chemical extracts (figure A2, table S2). In the extracts of egg surface compounds, we could observe between 21 (*A. senilis* and *L. niger)* and 31 (*C. aethiops*) peaks containing hydrocarbons that were consistently present in samples of the same species (figure A2, table S2). Profiles of eggs appear to contain a higher diversity of methyl-alkanes compared to linear alkanes. In *T. darioi, L. niger* and *F. fusca* egg samples, we also observed a small number of alkenes. The chemical profile of L1 larvae appears to have a lower total amount of hydrocarbons compared to eggs (figure A3) as well as a smaller diversity of compounds (figure A2, table S2). We found between 5 (in *L. niger* and *F. fusca*) and 9 (in *C. aethiops*) peaks containing hydrocarbons with a majority of linear alkanes and a lower number of methyl-alkanes in almost all species. In *M. barbarus*, both families of compounds were present in similar numbers (table S2). We did not observe any alkenes among the surface hydrocarbons extracted from larvae. The most common compounds were *n*-C_23_, *n*-C_25_and *n*-C_27_(peaks 4, 21 and 45), which are present across all species in surface profiles of both eggs and larvae (figure 1, table S2). The alkane *n*-C_28_(peaks 59) was found in all egg samples. In almost all cases, compounds found in L1 larvae extracts were also present in eggs extracts (figure A2, table S2). The exceptions are n-C_21_(peak 1), found on *A. senilis* and *L. niger* larvae only, and a diMeC_24_(peak 15) found on *A. senilis* larvae but not eggs.

**Figure 1:**
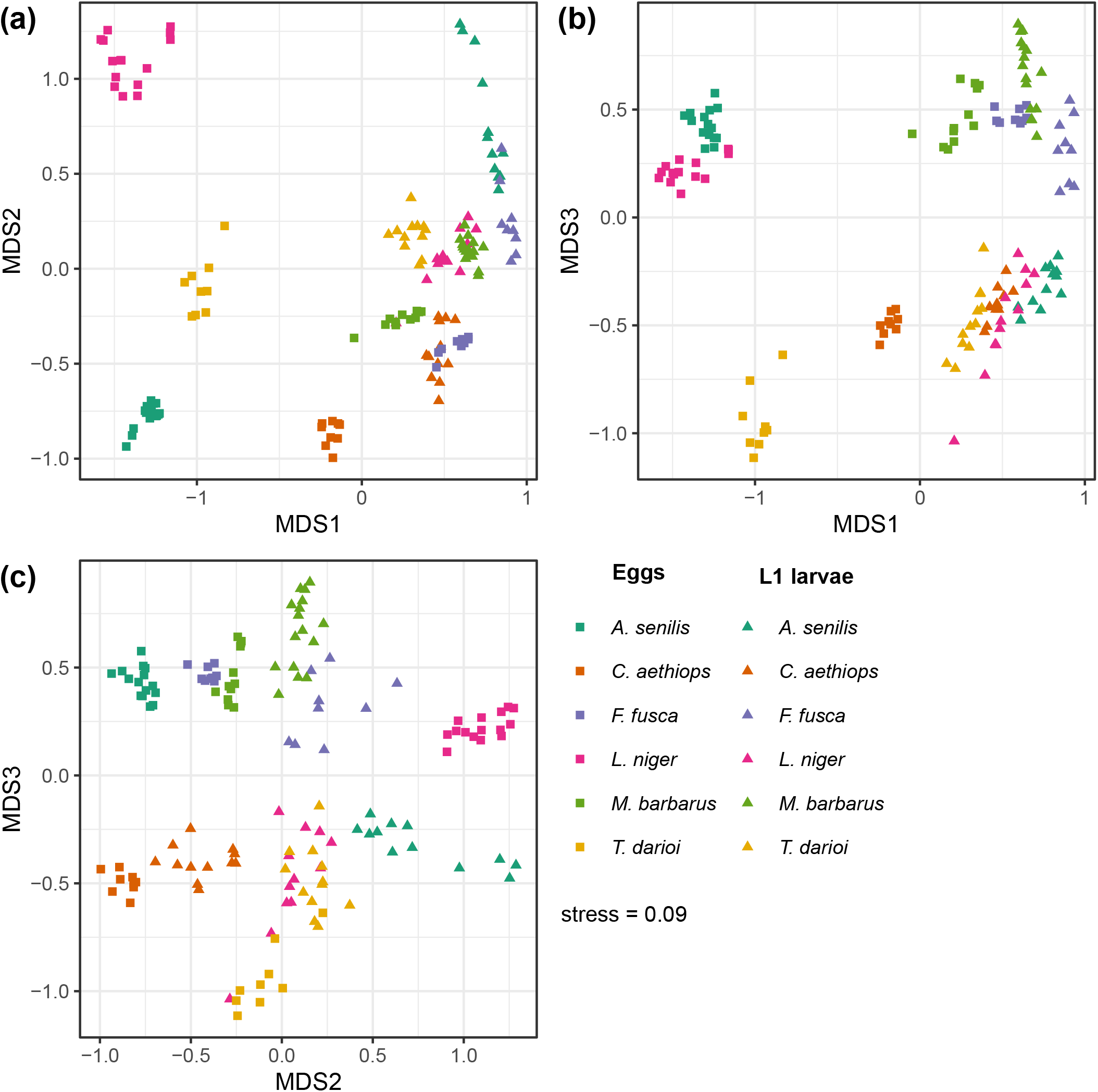
Chemical profiles of egg and L1 larvae. Scatterplots of non-metric multidimensional scaling of the area of hydrocarbons in surface extracts of eggs and L1 larvae with three output dimensions. **A)** Plot of first and second dimensions. B) Plot of first and third dimensions. C) Plot of second and third dimensions. Data points relative to egg samples are displayed with squares and L1 larvae with triangles. *A. sensilis* data point are plotted in dark green, *C. aethiops* in orange, *F. fusca* in violet, *L. niger* in magenta, *M. barbarus* in light green & *T. darioi* in yellow. Information on the origins of the sample extracted can be found in supplementary table S1. Non-ordinated chemical data is reported in supplementary table S2.

Principal component analyses indicate that there is a colony-specificity of surface hydrocarbons blends (figure A4, table A1). Using linear discriminant analyses, we observed that chemical profiles allowed the prediction of the colony of origin of the egg samples significantly better than by chance (permutation test, *P* ≤ 0.05, figure 2.a, Koul et al., 2018; Ojala & Garriga, 2010). The accuracy of prediction of the colony of origin was 100% for *L. niger, C. aethiops, F. fusca and M. barbarus* eggs. For *T. darioi* and *A. senilis* eggs, the prediction of the colony of origins was not totally accurate (88.89% and 93.33% respectively). In larvae samples, the hydrocarbon profiles allowed the identification of the colony of origin in *L. niger, C. aethiops, F. fusca* and *M. barbarus* (permutation test, *P* ≤ 0.05, figure 2.a). However, unlike for egg samples, the accuracy of prediction of the colony of origin was 100% only for *C. aethiops* and *F. fusca*. Regarding *M. barbarus* and *L. niger* L1 larvae, the predictions were imperfect (50.00% and 58.33% respectively). For *T. darioi* and *A. senilis* L1 samples, the prediction of the colony of origin was inaccurate (33.34% and 25.00% respectively) and not different from random (permutation test, *P* > 0.05, figure 2.a). Replication of this analysis with an NMDS ordination gave similar results, although PCA ordination appears to perform better (table A2).

**Figure 2:**
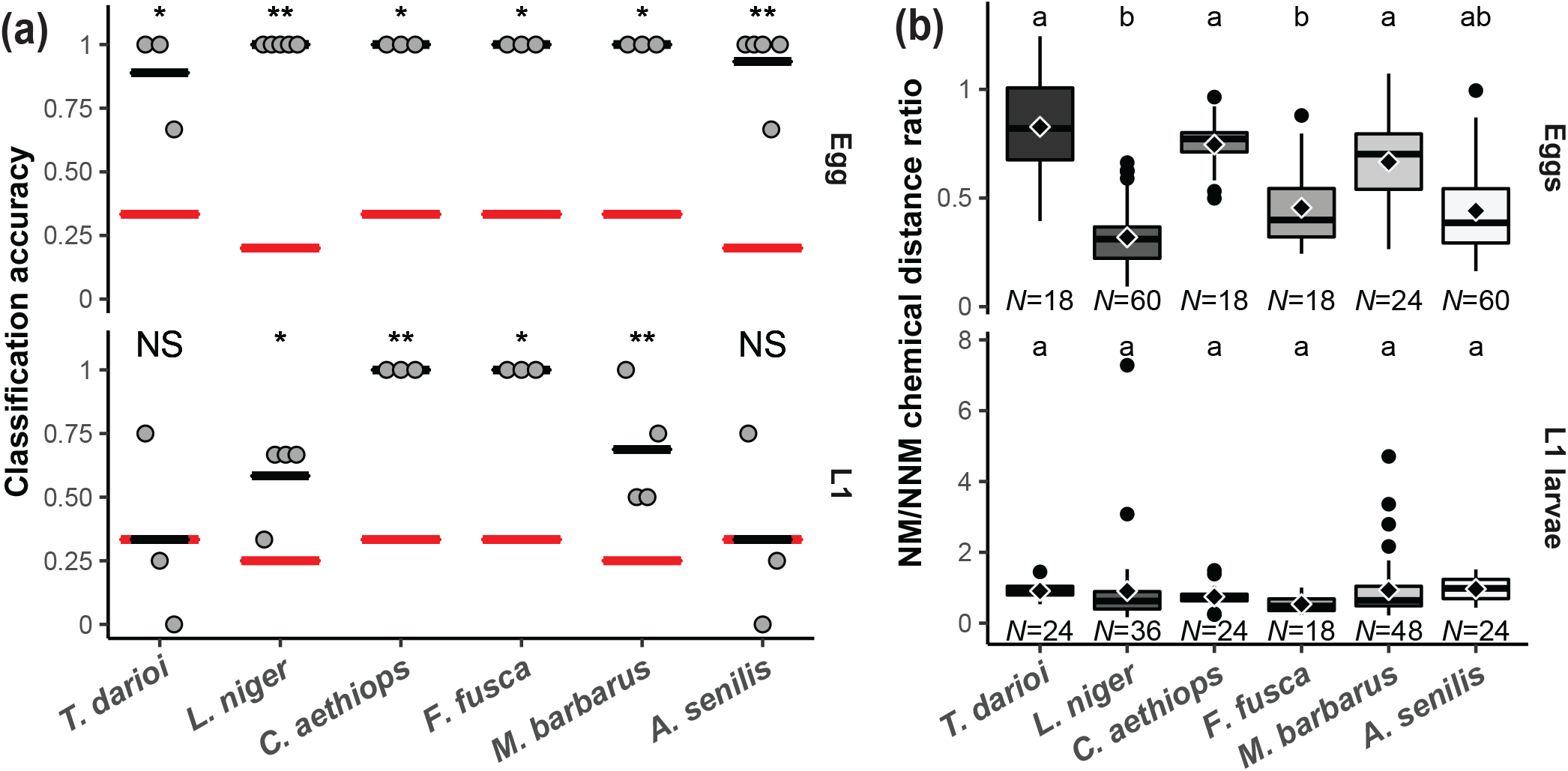
Colony specific hydrocarbon signature of ant early brood. Results of the analysis on chemical extracts of eggs and L1 larvae. Details on the origin of the samples used are displayed in supplementary table S1 **A)** Precisions of the linear discriminant analysis for each colony in each sample types performed from the principal components that had an F-score superior or equal to 0.01. The black narrower lines represent the mean precision for each sample type. The red wider lines represent accuracy expected from random choices. Significance of the difference of mean precisions compared to a random accuracy was computed with a permutation test. NS: p ≥ 0.05; *: *P* ≤ 0.05; **: *P* ≤ 0.01; ***: *P* ≤ 0.001. **B)** Ratios of the chemical Euclidean distances between nestmate and non-nestmate measured with the global-centroid method from the principal components that had an F-score superior or equal to 0.01. The sample size refers to number of distances measured between one sample and the samples from one of the others colonies of matching species and brood type. Black dots represent outlier values that are 1.5 times outside the interquartile range. Letters represent groups of statistical similarity in each sample type (LMM; Type II ANOVA; *P* ≤ 0.05).

To compare the difference between colony hydrocarbon profiles across species, we normalized the nestmate chemical distances relative to the non-nestmate distances in each species (figure2.b). The difference in colony signatures are similar for larvae and for eggs in most species. However, in *L. niger* and *F. fusca* eggs, the differences in colony signatures are larger compared to *T. darioi, C. aethiops* and *M. barbarus* nestmate to non-nestmate distances (LMM, *P* ≤ 0.05, Type II ANOVA; table A3). Consistently with our analysis of the existence of a colony signature in the chemical profiles of eggs, the large majority of ratios between nestmate and non-nestmate eggs chemical distances are inferior to one (*i.e*. distance between nestmates is smaller than between non-nestmates). For larvae, cases of ratios superior to one (*i.e*. distance between nestmates is greater than between non-nestmates) appear more frequently, which is consistent with our observations that colony signatures are less clear for L1 larvae.

### Brood discrimination by ant workers

From the results of our chemical analyses, we would predict that ant workers are able to discriminate between homo-specific and hetero-specific brood. The discrimination between nestmate and non-nestmate would be possible for eggs but more difficult for L1 larvae, especially in *A. senilis* and *T. darioi*. Using behavioural assays, we measured the number of brood items retrieved by workers (figure 3.a) as well as the time they spent antennating the brood (figure 3.b).

**Figure 3.**
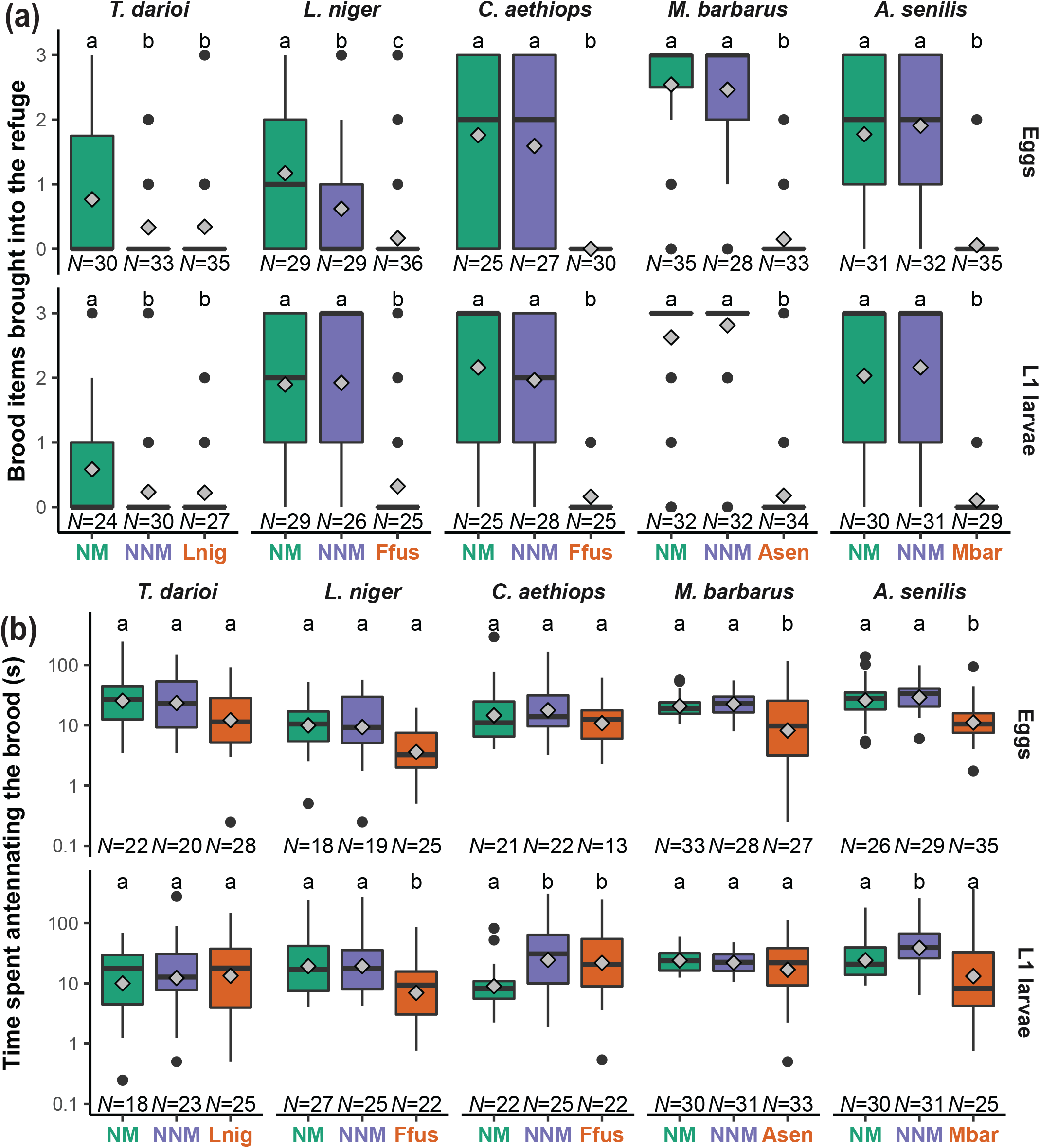
Worker behaviour towards early brood. Results of behavioural experiments performed on groups of 6 workers that were presented with 3 eggs or L1 larvae that were either nestmate (NM), non-nestmate (NNM) or hetero-specific (Asen: *A. senilis*, Mbar: *M. barbarus*, Lnig: *L. niger*, Ffus: *F. fusca*). **(a)** Boxplots of the number of brood items brought into the refuge by workers in the behavioural trials. The sample size refers to the number of worker groups that were active during a given trial (the total number of worker groups tested is reported in supplementary table S1). Bottom, middle and top horizontal lines of the box represent the first, the second and the third quartile, respectively. Horizontal lines represent the rest of the data range; black dots represent possible outlier values (1.5 times outside the interquartile range). **(b)** Boxplots of the total time spent by workers antennating brood during the trials. Diamonds represent the means. The sample size refers to the number of worker groups that were active and displayed antennation behaviour during a given trial (the total number of worker groups tested is reported in supplementary table S1). Letters show groups of statistical similarity in each species (LMM; Type II ANOVA; *P* ≤ 0.05). Boxplots represent data as in (a).

For *T. darioi*, nestmate eggs were retrieved significantly more frequently compared to hetero-specific items (GLMM, *P* ≤ 0.05, Type II ANOVA; table A4). We observed no differences in the number of non-nestmate and hetero-specific eggs retrieved by *T. darioi* workers. *L. niger* workers brought significantly more nestmate eggs into the refuge compared to non-nestmate and hetero-specific eggs (GLMM, *P* ≤ 0.05, Type II ANOVA; table A4). The number of non-nestmate eggs retrieved by *L. niger* workers was higher than the number of hetero-specific ones. The results for *A. senilis, C. aethiops, L. niger* and *M. barbarus* assays were similar: workers transported significantly more nestmate and non-nestmate eggs than hetero-specific ones into the refuge (GLMM, *P* ≤ 0.05, Type II ANOVA; Table A4). There was no significant difference between the number of nestmate and non-nestmate eggs retrieved by workers.

Regarding L1 larvae, *T. darioi* workers retrieved significantly more nestmate L1 larvae than non-nestmate and hetero-specific ones. In fact, *T. darioi* workers retrieved almost no non-nestmate or hetero-specific larvae. Consequently, there were no differences in the number of non-nestmate and hetero-specific larvae retrieved by *T. darioi* workers. Observations for *L. niger, A. senilis, C. aethiops*, and *M. barbarus* L1 larvae trials were similar between each other. The number of nestmate and non-nestmate L1 larvae transported into the refuge by workers were similar and significantly higher than the number of hetero-specific L1 larvae. Overall, the results of the behavioural assays show that ant workers are able to discriminate between homo-specific and hetero-specific eggs and L1 larvae. Furthermore, we observed that *L. niger* and *T. darioi* discriminate between nestmate and non-nestmate eggs and only *T. darioi* workers discriminate between nestmate and non-nestmate L1 larvae.

Antennation allows ants to use their chemical and mechanical sensors to explore items. A longer antennation time is a sign of a higher interest or more complex identification of the item. *A. senilis* and *M. barbarus* workers spent significantly more time antennating nestmate and non-nestmate eggs compared to hetero-specific eggs (LMM, *P* ≤ 0.05, Type II ANOVA; table A5). *L. niger* workers antennated for a significantly longer time nestmate and non-nestmate L1 larvae when compared to hetero-specific ones (LMM, *P* ≤ 0.05, Type II ANOVA; table A5). For *C. aethiops*, antennation times were significantly shorter when comparing nestmate to non-nestmate and hetero-specific L1 larvae (LMM, *P* ≤ 0.05, Type II ANOVA; table A5). Finally, *A. senilis* workers spent less time antennating nestmate and hetero-specific L1 larvae compared to non-nestmate larvae (LMM, *P* ≤ 0.05, Type II ANOVA; table A5).

Overall, our behavioural trials show that ant workers discriminate between brood items from their colony and hetero-specific ones. However, discrimination between nestmate and homo-specific non-nestmate brood is clearly evident only in *L. niger* and *T. darioi*.

## Discussion

Our chemical analyses and behavioural experiments allow a better understanding of species and colony-level chemical cues in the early brood stages of derived ant species as well as the discriminatory behaviour that could depend on those cues. The number of chemical cues observed is smaller in first instar larvae compared to eggs in all species studied. It seems to be the case for the diversity of surface hydrocarbons. However, we can’t rule out that our method of analysis, for which quantity of hydrocarbons appears limiting, was not sensitive enough to detect the full diversity hydrocarbons on larvae’s surface. Nevertheless, the difference in surface hydrocarbon quantity supports the hypothesis that when larvae hatch from the egg the hydrocarbons are not transferred from the egg’s chorion to the larval cuticle, or at least they are in very minute amounts. If so, L1 larvae would have to synthesize *de novo* their surface hydrocarbons. Transfer of hydrocarbon from workers might also be a way for larva to acquire the colony signature.

The hydrocarbons observed on the surface of eggs and L1 larvae are of a similar nature to those found in adults that were detected across a wide range of Hymenoptera species (Provost et al., 1994; van Zweden & d’Ettorre, 2010). As such, they should be detected by the sensory systems of most, if not all, ant species (Sharma et al., 2015). Our chemical analysis clearly showed that the surface hydrocarbons of eggs and L1 larvae are different among species. These inter-specific differences are consistent with our observation that ant workers discriminate both eggs and larvae of their species from brood of a different species in all our behavioural trials. This is also consistent with what has been observed for eggs in some *Formica* species (Chernenko et al., 2011; Schultner & Pulliainen, 2020).

Are ants able to recognize the colony of origin of conspecific eggs? We observed colony-specific blend of hydrocarbons on eggs, suggesting that the display of colony cues on eggs is a trait present across the three ant subfamilies we studied, which derived more than a 100 million years ago (Moreau et al., 2006). This is consistent with observations in seven *Formica* species (Helanterä & d’Ettorre, 2015). Despite the presence of colony-specific cues, only *T. darioi* and *L. niger* workers discriminated against non-nestmate eggs in our behavioural trials. Data from the literature show that *F. fusca* workers and larvae discriminate against non-nestmate eggs (Helanterä & Sundström, 2007; Pulliainen et al., 2019). Interestingly, our results showed that discrimination against non-nestmate eggs is not consistently corelated with larger differences between nestmate and non-nestmate odours. This indicates that non-nestmate discrimination could also rely on a more accurate recognition by workers of the cues displayed on the brood or on variation in the acceptance threshold of workers.

Can workers recognize nestmate first instar larvae? Our chemical analysis and behavioural trials with L1 larvae draw a less clear picture than for eggs. Data in the literature are also scant. Larvae from both Formicinae species we studied (*L. niger* and *C. aethiops*) and those from *M. barbarus* (Myrmicinae) display a colony-specific chemical signature. However, these signatures did not allow for reliable identification of the colony of origin by our analytical tools in two species (*M. barbarus* and *L. niger*). We could not demonstrate the presence of a colony signature in the surface hydrocarbons of *T. darioi* (Dolichoderinae) and *A. senilis* (Myrmicinae) larvae. Surprisingly, *T. darioi* workers were the only ones able to discriminate between nestmate and non-nestmate larvae, which indicates that *T. darioi* larvae display enough cues for colony-level recognition. This means that *T. darioi* workers either use chemical cues that our method of analysis could not detect or use non-chemical cues. However, to our knowledge, the literature does not support the hypothesis that workers use non-chemical cues (*e.g*. visual or auditory) for nestmate larvae recognition (Schultner & Pulliainen, 2020). As such, the hypothesis that *T. darioi* first instar larvae display a colony specific odour remains the most plausible.

Our experimental setup required compromises to allow testing multiple species in a comparable way. Trials were performed on small groups of individuals compared to the size of ant colonies in nature. However, we used refuges that were previously stored in the colony of origin to allow these refuges to bear the colony’s odour. We also made sure that ant groups were accepting the refuge as a suitable brood storage by selecting groups that displayed a brood retrieval behaviour during the acclimation stage. The worker groups we used could be considered as recently queen-less. Nevertheless, the workers should be able to sense the queen presence from the three pieces of brood they had in their refuge. As such, we are confident that the behaviour of the workers in our set-up was not altered in a way that would impair our conclusion.

We observed *A. senilis* and *C. aethiops* workers behaving differently when facing nestmate larvae compared to non-nestmate larvae (*i.e*. different antennation durations). Is this an indication that they are able to recognize nestmate L1 larvae from non-nestmate larvae? On *C. aethiops* L1 larvae, we could detect a colony-level chemical signature. We could not reliably do so on *A. senilis* first instar larvae, but neither could we on *T. darioi* larvae despite the clear behavioural evidences that they do display a colony signature. Given the lower overall quantity of surface hydrocarbons on L1 larvae compared to eggs, the chemical cues displayed might challenge the olfactory detection system of ant workers and the presence of non-nestmate cues might appear ambiguous to them. The long antennation time observed would then be a sign of the ant’s difficulty to recognize the signature. Similar hesitation has been observed for recognition of ambiguous colony cues on adults (Nascimento et al., 2013).

Taken together, our observations allow us to confidently state that workers recognize and favour nestmate first instar larvae only for *T. darioi*. In the other species, discrimination is clear only towards hetero-specific larvae. Discrimination against non-nestmate eggs, doesn’t implies favouring nestmate first instar larvae. These differences across stages in non-nestmate discrimination probably arose from the differences in the quality and the diversity of the chemical cues displayed as the surface of the brood. Unlike eggs, larvae likely have to synthesize the chemical cues they display from the first day after emergence. It is also possible that the difference in discriminatory behaviour of *L. niger* towards eggs and L1 larvae are linked to a risk-reward trade-off between these two brood stages. L1 larvae need a shorter time, hence less resources, to become adult workers compared to eggs.

Our observations and those from the literature support the hypothesis that egg surface hydrocarbons display sufficient information for ant workers to discriminate nestmate from non-nestmate eggs across the most derived clades of the ants’ phylogenetic tree. The predominance of non-nestmate eggs discrimination in the majority of the ant species studied calls for further work, on additional ant species, to test evolutionary hypotheses on conspecific non-nestmates discrimination in ants.

The three ant species that efficiently discriminate against non-nestmate eggs belong to genera prone to social parasitism. Indeed, *L. niger* is host to various social parasites from the *Lasius* genus (Buschinger, 2009) and the *Tapinoma* genus is known to be subject to parasitism by *Bothriomyrmex* species (Buschinger, 2009; Lenoir et al., 2001). Furthermore, host species of the *Formica* genus also discriminate against non-nestmate eggs (Chernenko et al., 2011). Our results, and those from the literature, are thus in accordance with the hypothesis that the arms race between social parasites and host species led workers from host species to set an adaptatively less permissive acceptance threshold regarding divergence from the colony signature on brood, thus discriminating against non-nestmates (Pulliainen et al., 2019). The parasites trying to get themselves recognized as nestmates induce a more strict discrimination of eggs as a species level adaptation in hosts (Grüter et al., 2018).

Discrimination can lead to costly errors (Reeve, 1989; Rossi et al., 2018). Accordingly, the three species we studied, which are not subject to an arms race with social parasites, do not discriminate against non-nestmate brood. Brood adoption appears less risky in those non-host species while recognition errors (discarding of nestmate brood) represent a potential loss to the colony’s fitness. This would explain the reduction or disappearance of the discriminatory behaviour against non-nestmate eggs. Identification of first instar larvae, which do not display as many chemical cues as eggs, appears a more challenging task, which prevents a stricter non-nestmate discrimination in most species, even parasitized ones. Overall, our results are in accordance with the hypothesis that differences in selective pressures induced by social parasites are linked with differences in the discrimination against non-nestmate eggs in the context of brood retrieval between host and non-host species.

Given the relative artificial nature of our experimental set-up, we can however not rule out that the recently queen-less workers would be overall more prone to retrieve brood. As such, our experimental set-up would then have induced a higher non-nestmate brood retrieval, without masking the difference in behaviour between host and non-host species. As observed here in the case non-host ant species, there are other species of ants and social insects in general, that do not discriminate against non-nestmates, or non-kin, even though theory would predict them to do so (Blatrix & Jaisson, 2002; de Gasperin et al., 2021; Friend & Bourke, 2012; Helanterä et al., 2007; Kikuchi et al., 2007; Mora-Kepfer, 2014). Outside social insects, bird or mammals can either be kin-discriminative or not in their altruistic behaviour depending on the species. A possible explanation is the fact that group members are usually highly related and errors cost more than providing resources to less related offspring (Duncan et al., 2019). Overall, this suggests that discrimination strategies often result from trade-offs and depends on organisms’ life-history and ecology.

## Appendices

**Table A1:**
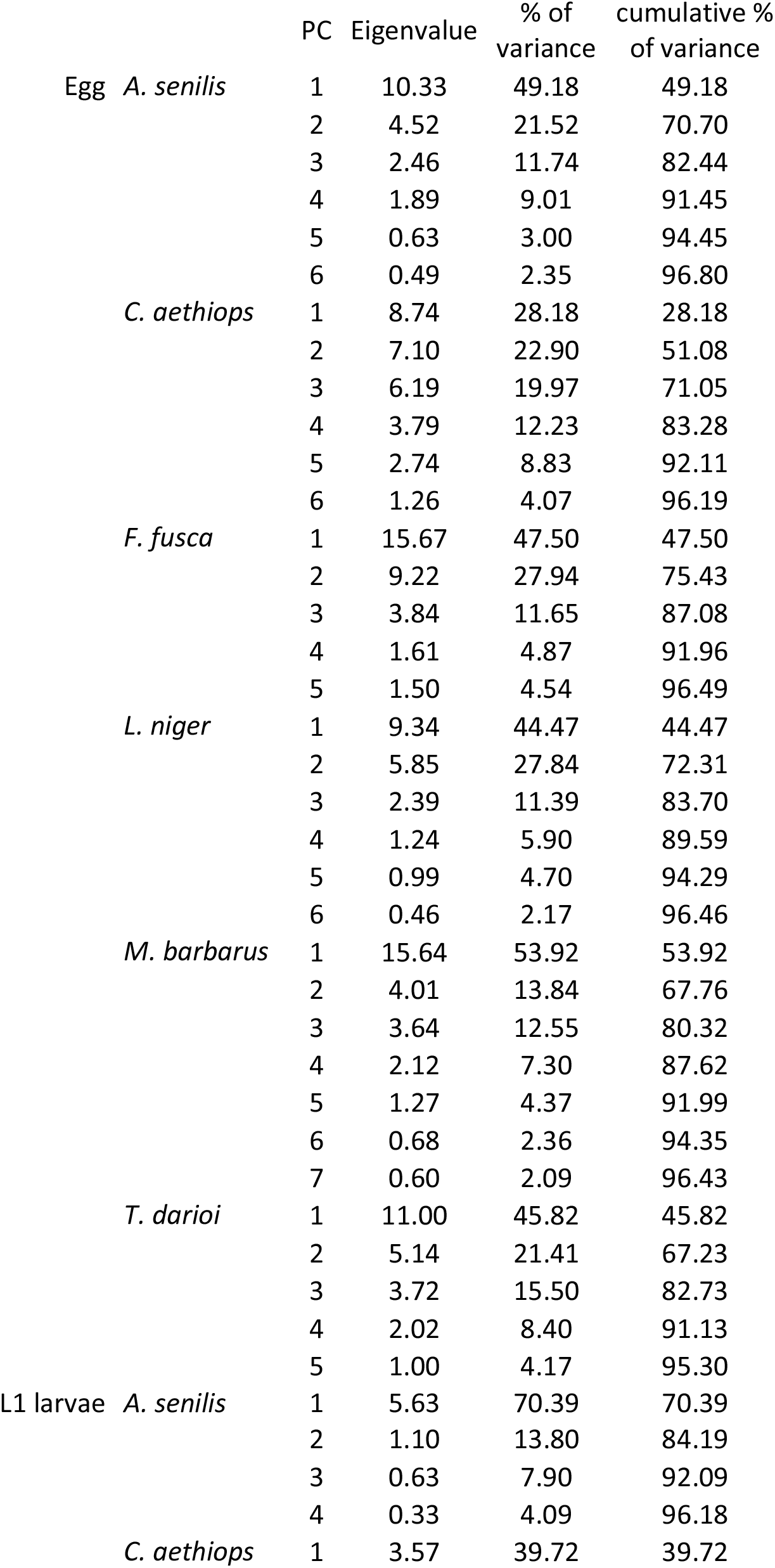
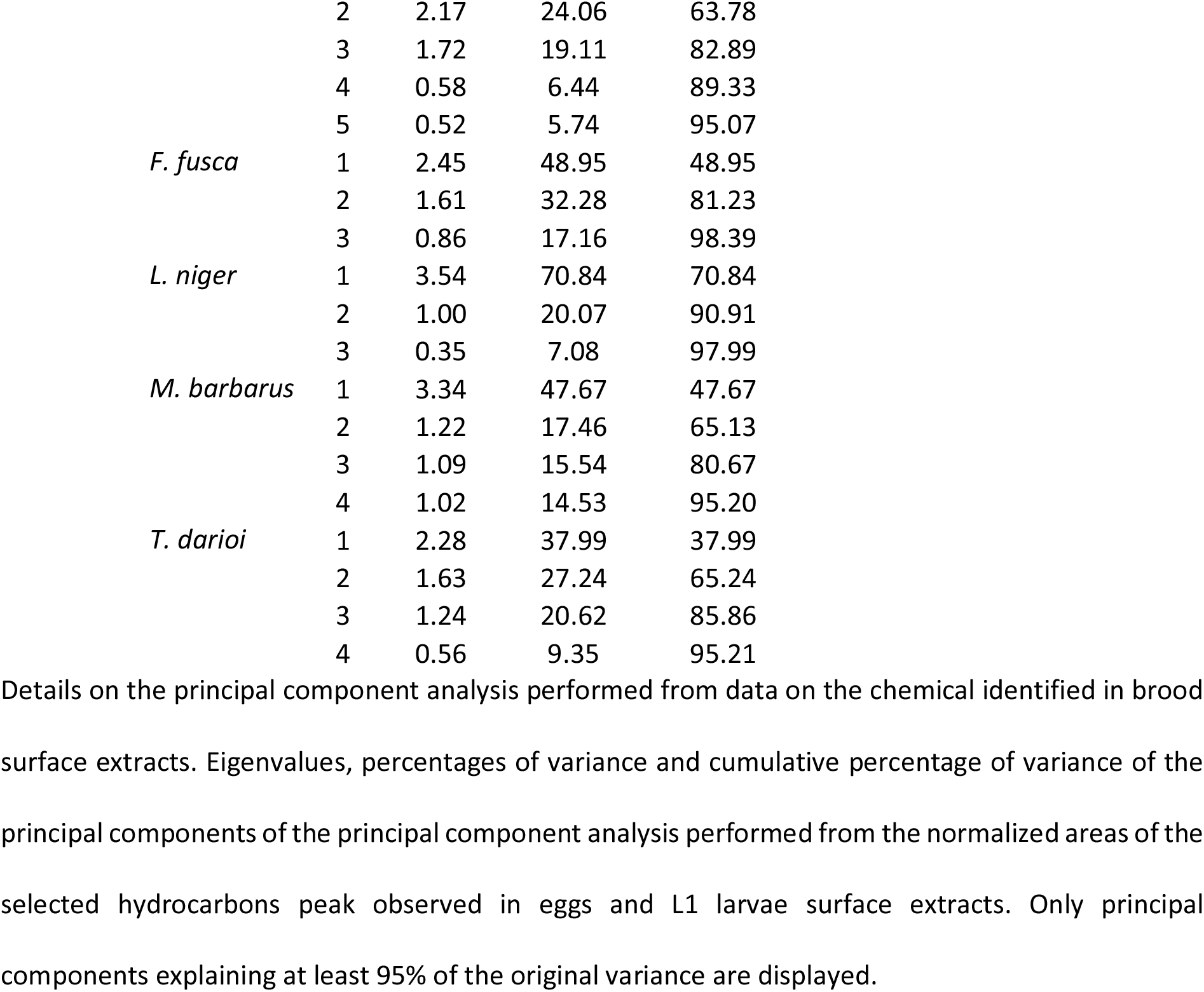
Percentage of variance explained by the dimensions of the principal component analyses.

|  |  | PC | Eigenvalue | % of variance | cumulative % of variance |
| --- | --- | --- | --- | --- | --- |
| Egg | *A. senilis* | 1 | 10.33 | 49.18 | 49.18 |
|  |  | 2 | 4.52 | 21.52 | 70.70 |
|  |  | 3 | 2.46 | 11.74 | 82.44 |
|  |  | 4 | 1.89 | 9.01 | 91.45 |
|  |  | 5 | 0.63 | 3.00 | 94.45 |
|  |  | 6 | 0.49 | 2.35 | 96.80 |
|  | *C. aethiops* | 1 | 8.74 | 28.18 | 28.18 |
|  |  | 2 | 7.10 | 22.90 | 51.08 |
|  |  | 3 | 6.19 | 19.97 | 71.05 |
|  |  | 4 | 3.79 | 12.23 | 83.28 |
|  |  | 5 | 2.74 | 8.83 | 92.11 |
|  |  | 6 | 1.26 | 4.07 | 96.19 |
|  | *F. fusca* | 1 | 15.67 | 47.50 | 47.50 |
|  |  | 2 | 9.22 | 27.94 | 75.43 |
|  |  | 3 | 3.84 | 11.65 | 87.08 |
|  |  | 4 | 1.61 | 4.87 | 91.96 |
|  |  | 5 | 1.50 | 4.54 | 96.49 |
|  | *L. niger* | 1 | 9.34 | 44.47 | 44.47 |
|  |  | 2 | 5.85 | 27.84 | 72.31 |
|  |  | 3 | 2.39 | 11.39 | 83.70 |
|  |  | 4 | 1.24 | 5.90 | 89.59 |
|  |  | 5 | 0.99 | 4.70 | 94.29 |
|  |  | 6 | 0.46 | 2.17 | 96.46 |
|  | *M. barbarus* | 1 | 15.64 | 53.92 | 53.92 |
|  |  | 2 | 4.01 | 13.84 | 67.76 |
|  |  | 3 | 3.64 | 12.55 | 80.32 |
|  |  | 4 | 2.12 | 7.30 | 87.62 |
|  |  | 5 | 1.27 | 4.37 | 91.99 |
|  |  | 6 | 0.68 | 2.36 | 94.35 |
|  |  | 7 | 0.60 | 2.09 | 96.43 |
|  | *T. darioi* | 1 | 11.00 | 45.82 | 45.82 |
|  |  | 2 | 5.14 | 21.41 | 67.23 |
|  |  | 3 | 3.72 | 15.50 | 82.73 |
|  |  | 4 | 2.02 | 8.40 | 91.13 |
|  |  | 5 | 1.00 | 4.17 | 95.30 |
| L1 larvae | *A. senilis* | 1 | 5.63 | 70.39 | 70.39 |
|  |  | 2 | 1.10 | 13.80 | 84.19 |
|  |  | 3 | 0.63 | 7.90 | 92.09 |
|  |  | 4 | 0.33 | 4.09 | 96.18 |
|  | *C. aethiops* | 1 | 3.57 | 39.72 | 39.72 |

|  |  |  |  |  |
| --- | --- | --- | --- | --- |
|  | 2 | 2.17 | 24.06 | 63.78 |
|  | 3 | 1.72 | 19.11 | 82.89 |
|  | 4 | 0.58 | 6.44 | 89.33 |
|  | 5 | 0.52 | 5.74 | 95.07 |
| *F. fusca* | 1 | 2.45 | 48.95 | 48.95 |
|  | 2 | 1.61 | 32.28 | 81.23 |
|  | 3 | 0.86 | 17.16 | 98.39 |
| *L. niger* | 1 | 3.54 | 70.84 | 70.84 |
|  | 2 | 1.00 | 20.07 | 90.91 |
|  | 3 | 0.35 | 7.08 | 97.99 |
| *M. barbarus* | 1 | 3.34 | 47.67 | 47.67 |
|  | 2 | 1.22 | 17.46 | 65.13 |
|  | 3 | 1.09 | 15.54 | 80.67 |
|  | 4 | 1.02 | 14.53 | 95.20 |
| *T. darioi* | 1 | 2.28 | 37.99 | 37.99 |
|  | 2 | 1.63 | 27.24 | 65.24 |
|  | 3 | 1.24 | 20.62 | 85.86 |
|  | 4 | 0.56 | 9.35 | 95.21 |

Details on the principal component analysis performed from data on the chemical identified in brood surface extracts. Eigenvalues, percentages of variance and cumulative percentage of variance of the principal components of the principal component analysis performed from the normalized areas of the selected hydrocarbons peak observed in eggs and L1 larvae surface extracts. Only principal components explaining at least 95% of the original variance are displayed.

**Table A2:** Results of the statistical analysis of linear discriminant analysis

| Ordination method | Brood type | Species | Random accuracy | Classification accuracy | Adjusted *P* value |
| --- | --- | --- | --- | --- | --- |
| NMDS | Egg | *A. senilis* | 0.20 | 0.87 | **0.001** |
|  |  | *C. aethiops* | 0.33 | 0.89 | **0.018** |
|  |  | *F. fusca* | 0.33 | 1.00 | **0.013** |
|  |  | *L. niger* | 0.20 | 1.00 | **0.001** |
|  |  | *M. barbarus* | 0.33 | 0.75 | **0.018** |
|  |  | *T. darioi* | 0.33 | 0.33 | 0.166 |
|  | L1 larvae | *A. senilis* | 0.33 | 0.25 | 0.392 |
|  |  | *C. aethiops* | 0.33 | 0.92 | **0.001** |
|  |  | *F. fusca* | 0.33 | 0.67 | 0.150 |
|  |  | *L. niger* | 0.25 | 0.50 | **0.041** |
|  |  | *M. barbarus* | 0.25 | 0.44 | 0.114 |
|  |  | *T. darioi* | 0.33 | 0.42 | 0.216 |
| PCA | Egg | *A. senilis* | 0.20 | 0.93 | **0.001** |
|  |  | *C. aethiops* | 0.33 | 1.00 | **0.043** |
|  |  | *F. fusca* | 0.33 | 1.00 | **0.043** |
|  |  | *L. niger* | 0.20 | 1.00 | **0.001** |
|  |  | *M. barbarus* | 0.33 | 1.00 | **0.022** |
|  |  | *T. darioi* | 0.33 | 0.89 | **0.043** |
|  | L1 larvae | *A. senilis* | 0.33 | 0.25 | 0.812 |
|  |  | *C. aethiops* | 0.33 | 1.00 | **0.002** |
|  |  | *F. fusca* | 0.33 | 1.00 | **0.035** |
|  |  | *L. niger* | 0.25 | 0.58 | **0.039** |
|  |  | *M. barbarus* | 0.25 | 0.50 | **0.039** |
|  |  | *T. darioi* | 0.33 | 0.33 | 0.812 |

Details on the statistical analysis of the accuracy and results of permutation tests of the linear discriminant analysis with leave-one sample out cross-validation.

**Table A3:**
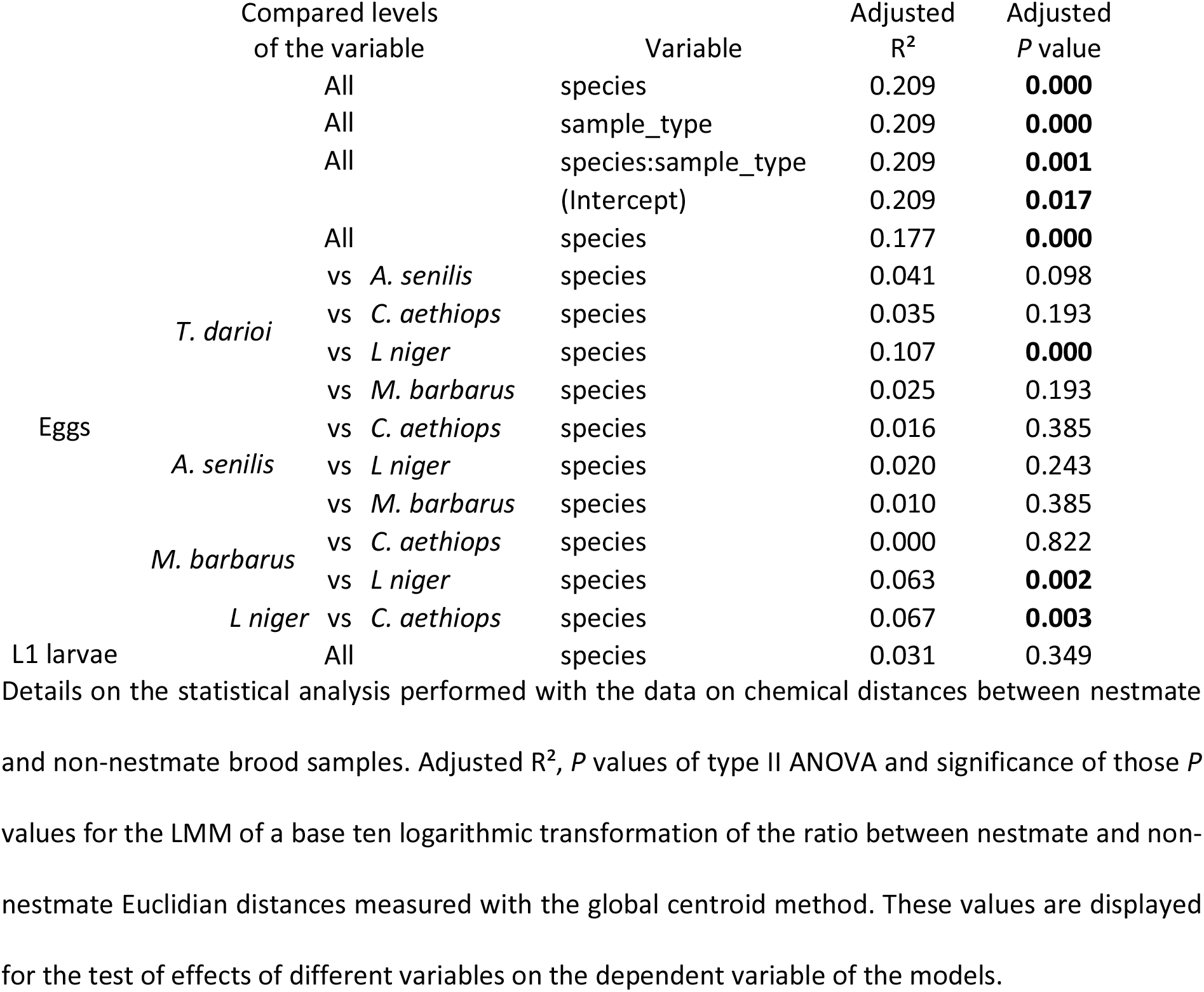
Results of the statistical analysis of nestmate / non-nestmate Euclidian distances

**Table A4:**
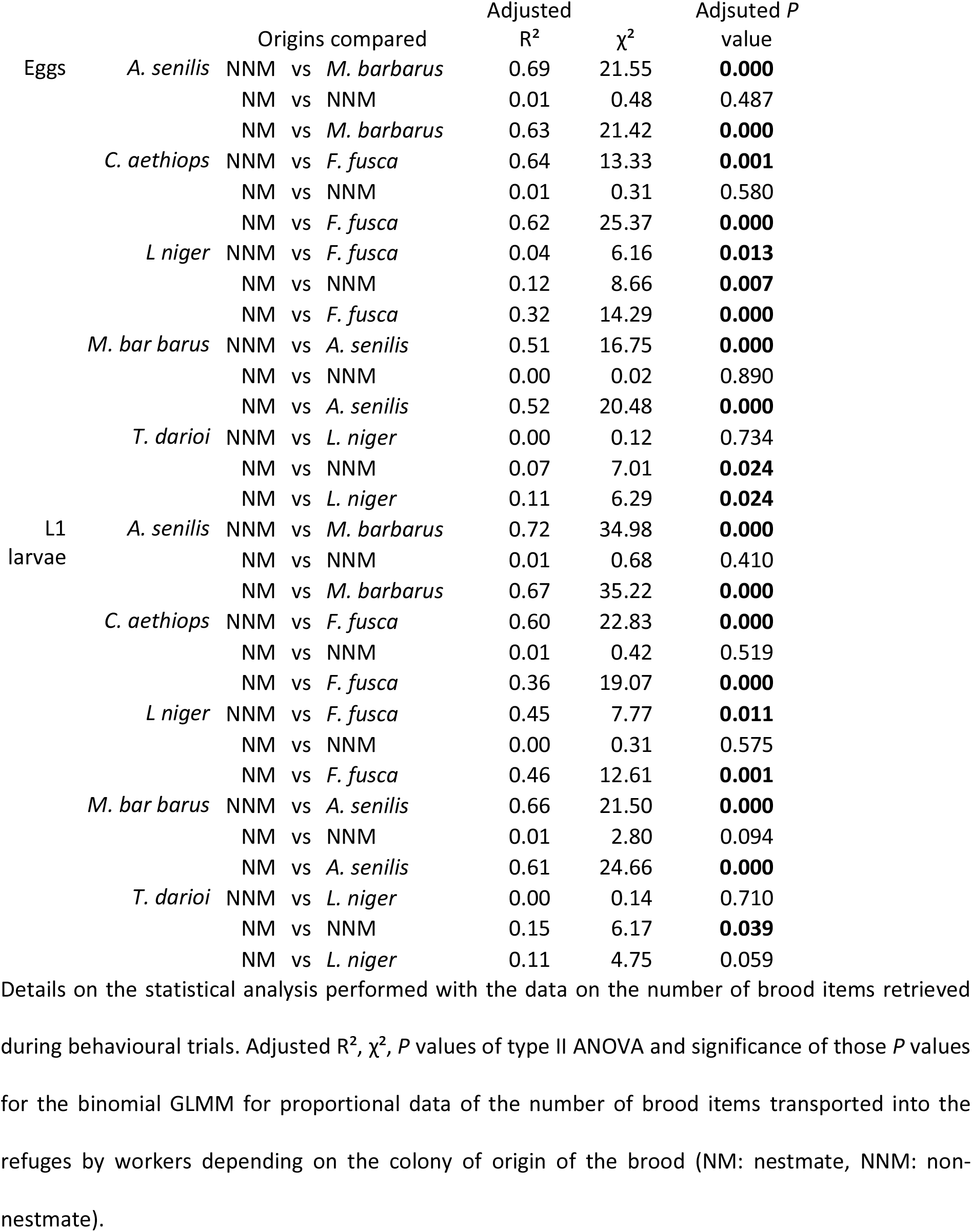
Results of the statistical analysis of the number of brood items transported into the refuge by workers

**Table A5:**
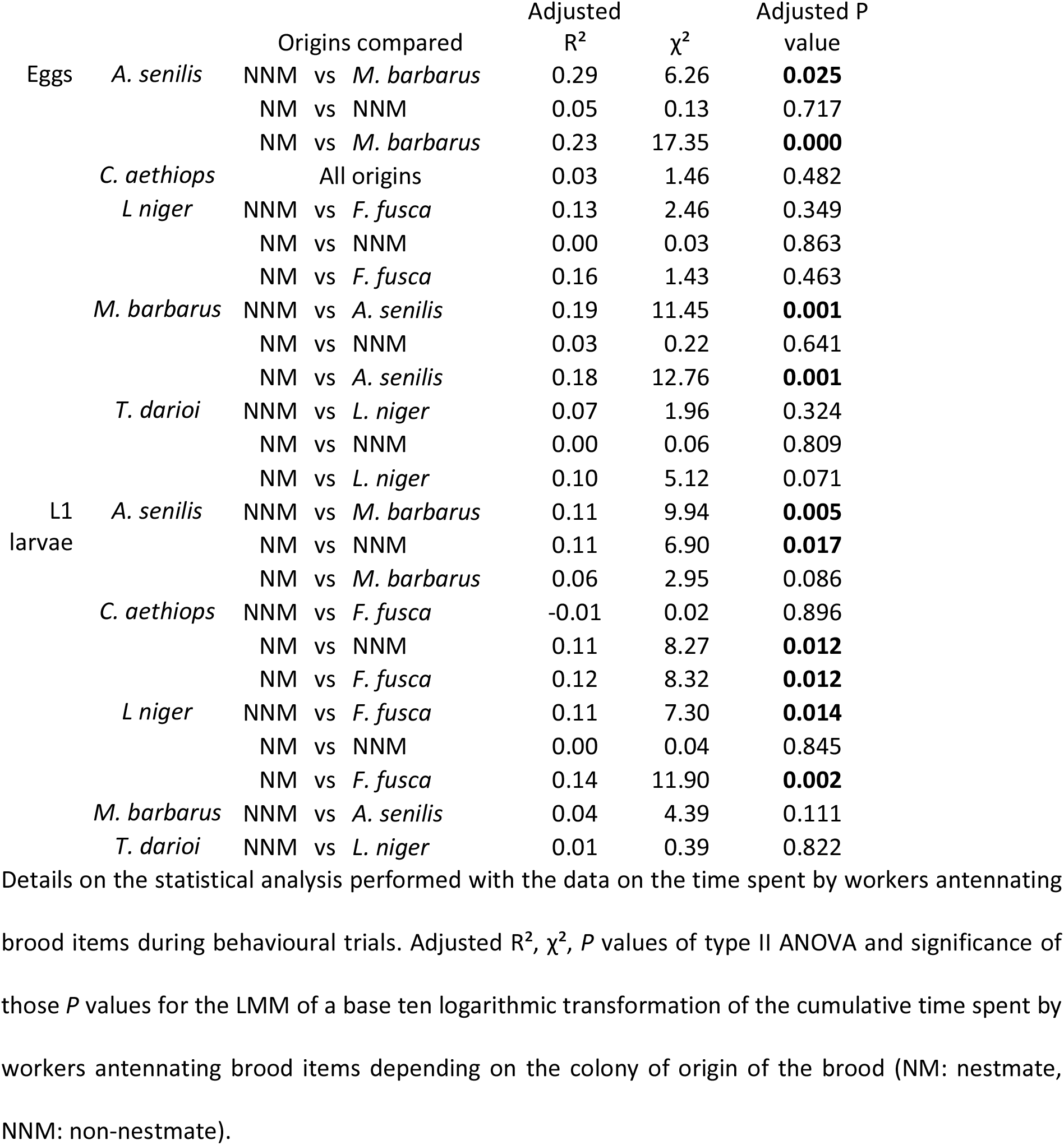
Results of the statistical analysis of the cumulative times spent by workers antennating brood items

## Figure legends

**Figure A1.**
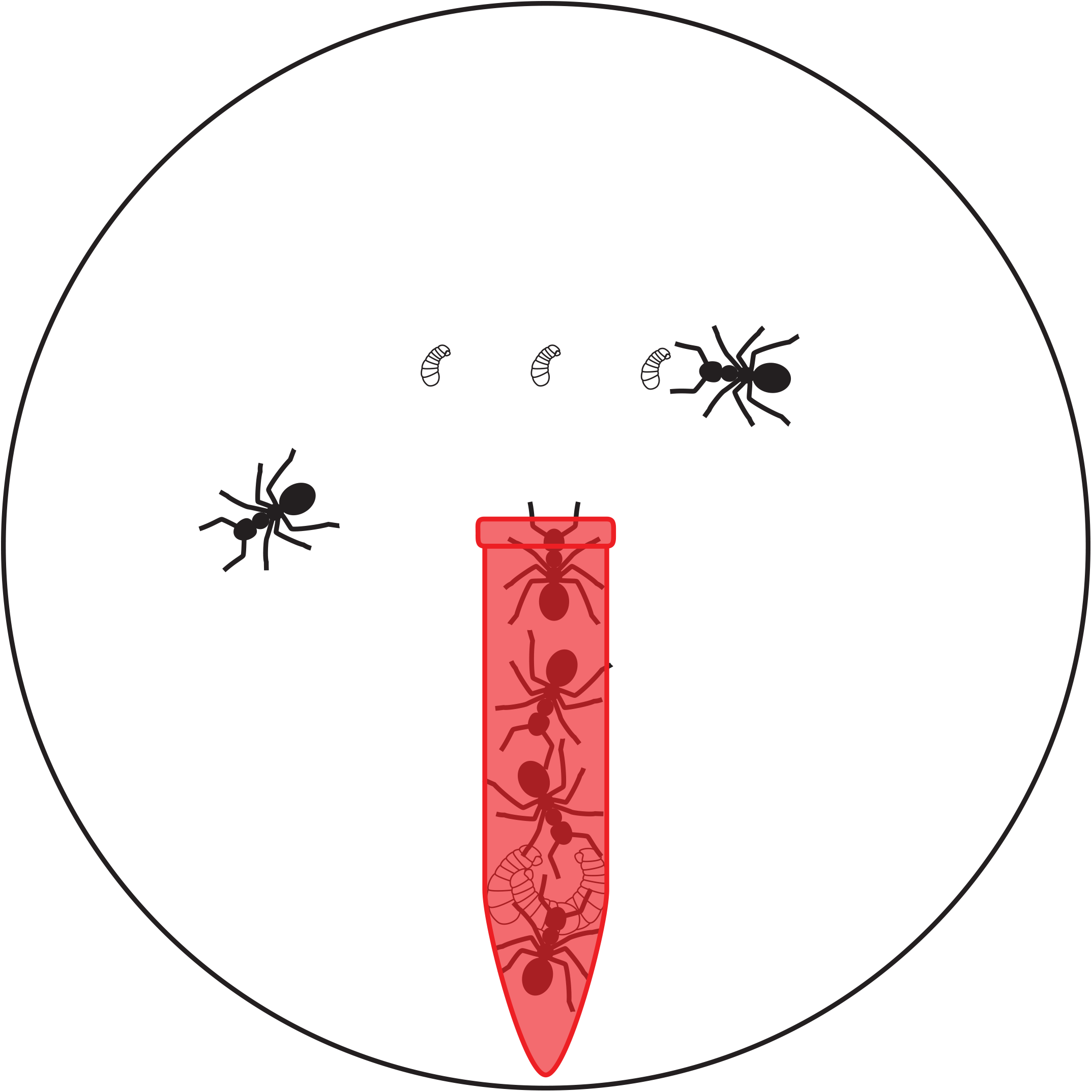
Disposition of the arenas of the behavioural assays. Diagram of the behavioural trial apparatus. Six workers (three from outside the nest and three from inside the nest) in an eight cm arena with Fluon®-coated walls and a filter paper as floor. The red tube is a refuge made of a red-coated 1.5mL Eppendorf tube that had spent at least twenty-four hours in the colony box. Inside the refuge, the three late-instar larvae were given to the worker 24h prior experimentation. Outside the refuge, the three L1 larvae (either nestmate, non-nestmate or hetero-specific) are the ones given to the workers during the trials.

**Figure A2.**
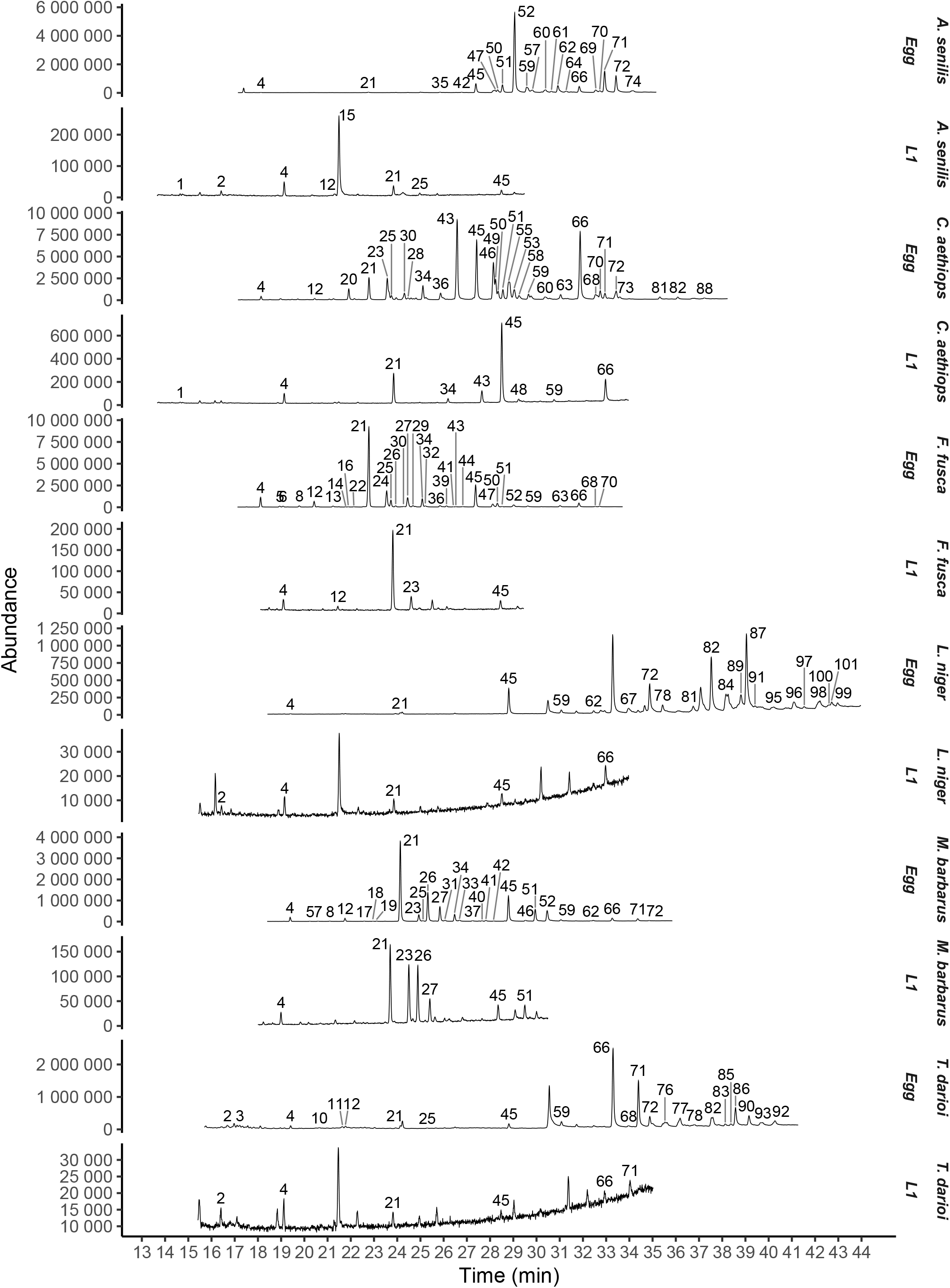
Brood items surface extracts. Representative chromatograms of surface extracts of *A senilis, C. aethiops, L. niger, M. barbarus* and *T. darioi* eggs and L1 larvae. Extracts were obtained from single eggs or larvae. Each peak with a number result from hydrocarbons that are found consistently across all samples of the same species and brood type (detailed in supplementary table S2).

**Figure A3.**
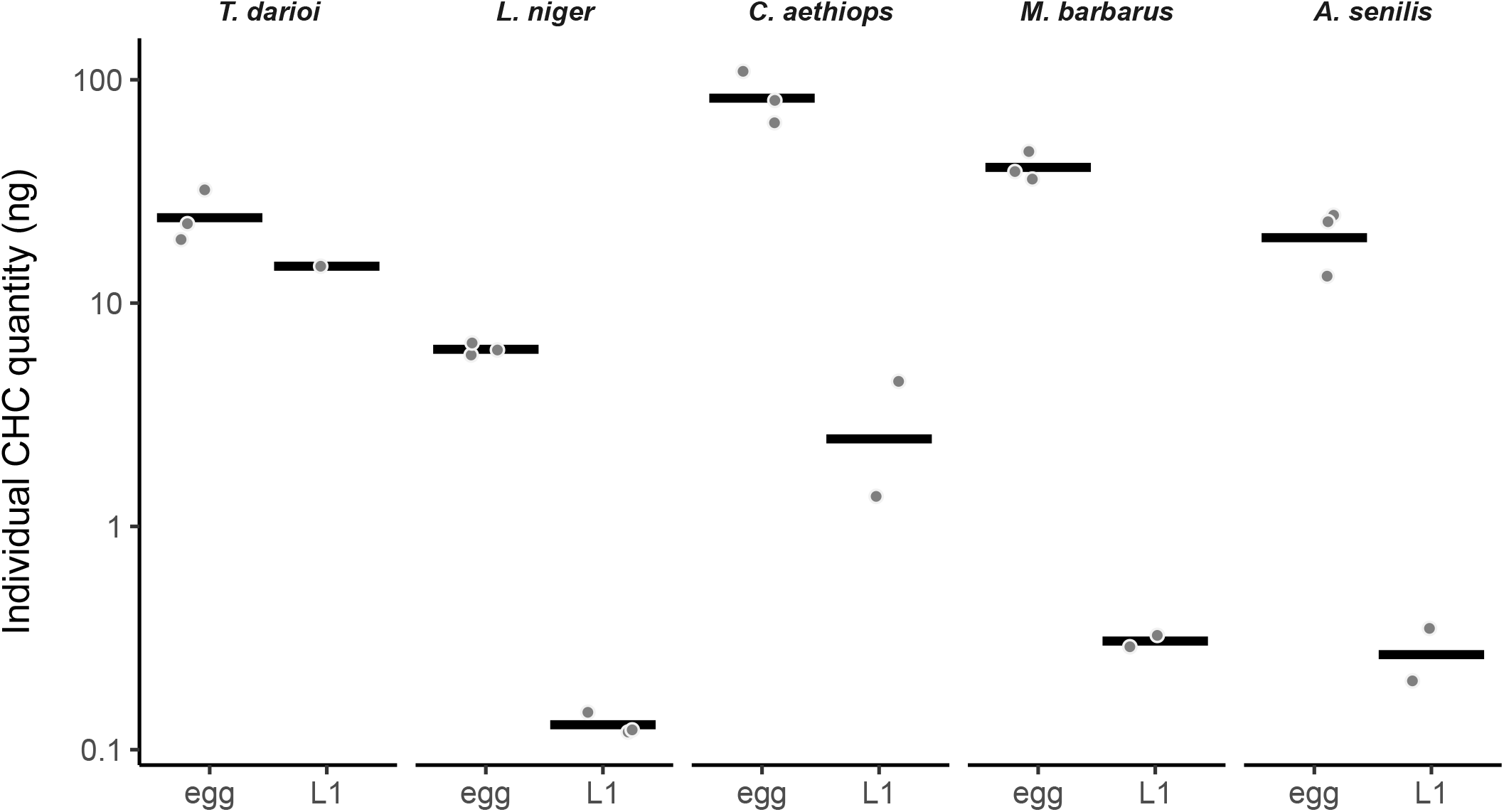
Quantity of surface hydrocarbons in eggs and L1 larvae extracts. Dot plots of the quantity (in ng) of hydrocarbons in surface extract of eggs and L1 larvae from *T. darioi, L. niger, C. aethiops, M. barbarus* and *A. senilis*. The black bar represents the mean for each species and sample type.

**Figure A4:**
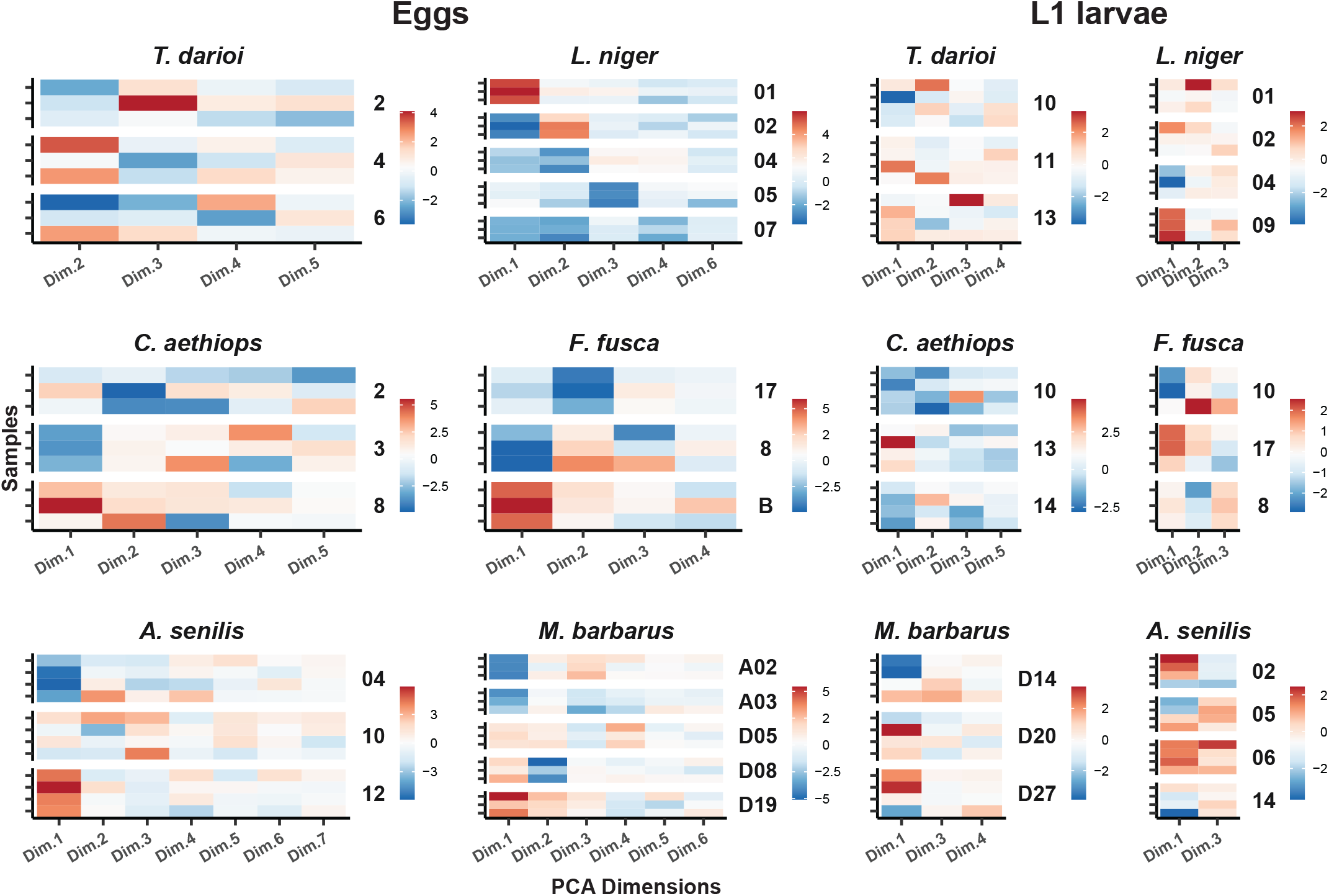
PCA dimensions heatmaps. Heatmaps of the principal components representing 95% of the initial variability of the normalized areas of the peaks obtained from surface extracts of eggs and L1 larvae. The values of the principal components are normalized relative to the highest absolute value observed for each principal component in each samples type. Each line is an individual sample. Samples from the same colony are grouped into the same square.

